# Compound FC-10696 Inhibits Egress and Spread of Marburg Virus

**DOI:** 10.1101/2021.01.15.426918

**Authors:** Ziying Han, Hong Ye, Jingjing Liang, Ariel Shepley-McTaggart, Jay E. Wrobel, Allen B. Reitz, Alison Whigham, Katrina N. Kavelish, Michael S. Saporito, Bruce D. Freedman, Olena Shtanko, Ronald N. Harty

## Abstract

Marburg virus (MARV) VP40 protein (mVP40) directs egress and spread of MARV, in part, by recruiting specific host WW-domain containing proteins via its conserved PPxY Late (L) domain motif to facilitate efficient virus-cell separation. We reported previously that small molecule compounds targeting the viral PPxY/host WW-domain interaction inhibited VP40-mediated egress and spread. Here, we report on the antiviral potency of novel compound FC-10696, which emerged from extensive structure activity relationship (SAR) of a previously described series of PPxY inhibitors. We show that FC-10696 inhibits egress of both mVP40 VLPs and egress and spread of authentic MARV from HeLa cells and primary human macrophages. Moreover, FC-10696 treated mice displayed delayed onset of weight loss, clinical signs, and significantly lower viral loads compared to controls, with 14% of animals surviving 21 days following a lethal MARV challenge. Thus, FC-10696 represents a first-in-class, host-oriented inhibitor effectively targeting late stages of the MARV lifecycle.

## Introduction

MARV is an emerging pathogen and potential bioterror agent that can cause severe hemorrhagic fever in humans and non-human primates (1). Currently, there are no approved vaccines or antiviral therapeutics to prevent or treat MARV infections. Development of novel and effective antiviral therapeutics against MARV and other members of the *Filoviridae* family are urgently needed.

MARV, like many other enveloped RNA viruses, relies on its matrix protein (mVP40) to direct and promote budding of infectious virions. We and others have demonstrated that mVP40 completes the budding process, in part, by using its highly conserved PPxY L-domain motif to hijack host proteins/pathways that then facilitate efficient virus-cell separation (2–16). One of the best characterized host interactors is WW-domain containing E3 ubiquitin ligase, Nedd4 (7, 16–18). Notably, we and others have shown that Nedd4 and Nedd4 family members physically interact with viral PPxY motifs via one or more of their WW-domains, and functionally interact with viral PPxY-containing proteins to enhance or promote budding of virus-like particles (VLPs) and live virus (7, 16–24). The highly conserved physical and functional nature of the PPxY motif in a wide array of RNA viruses, makes it an attractive target for the development of antivirals (2, 25–30). Indeed, compounds targeting the PPxY/WW-domain virus-host interaction would be predicted to dampen or reduce the ability of the virus to bud or pinch-off from infected cells, thus allowing an individual’s immune system more time to combat and clear the virus.

Previously, we described the identification and development of two novel series of small molecule compounds that significantly inhibited a VP40-Nedd4 interaction and PPxY-mediated egress of filovirus VP40 VLPs (26). We went on to show that our lead compounds significantly inhibited budding of a live VSV recombinant (VSV-M40) that we engineered to express the PPxY motif from Ebola virus (EBOV) VP40 in place of that from the VSV matrix (M) protein (26). Following extensive SAR and analog testing, we have now identified lead compound FC-10696, which we have shown is stable in human liver microsomes and possesses suitable ADME and PK properties for IP administration and testing in mice. Indeed, here we show that nanomolar concentrations of FC-10696 blocked budding of mVP40 VLPs. Moreover, we demonstrate that similar concentrations of FC-10696 also significantly inhibited egress and spread of live MARV in both HeLa cells and human monocyte derived macrophages (hMDM) with little to no cytotoxicity at the effective antiviral concentrations. Importantly, we show proof-of-concept *in vivo* efficacy of FC-10696 in a mouse challenge model of MARV disease. Indeed, FC-10696-treated animals exhibited delayed onset of weight loss, clinical signs, and disease progression compared to control animals, with 14% of animals from the FC-10696 treated group surviving a lethal MARV challenge up to 21 days post-infection. These findings represent the first proof-of-concept efficacy study for a novel host-oriented antiviral that has the potential for broad-spectrum activity against other PPxY containing viruses such as EBOV and Lassa virus (LASV), and provides proof-of-principle for further development of these first-in-class compounds targeting viral L-domain/host interactions as effective countermeasures to reduce virus budding and dissemination.

## Results

### ADME, PK, and anti-budding data for compound FC-10696

We improved drug disposition properties of a previously described series (26) to generate a novel small molecule FC-10696, a compound with vastly improved overall ADME and PK properties (Table 1), and with robust anti-budding activity against mVP40 VLPs (Table 1 and Fig. 1). Indeed, results from a representative mVP40 VLP budding assay highlight the dose-dependent decrease in mVP40 VLP egress from HEK239T cells treated with increasing concentrations of FC-10696 (Fig. 1, lanes 3-6) compared to that from a vehicle (DMSO) alone treated control (Fig. 1, lane 1). Previously described active compound **1** (FC-4005) at 1.0μM served as a positive control (Fig. 1, lane 2) (26). As expected, equivalent amounts of mVP40 were detected in cell lysates from all samples (Fig. 1, Cells, lanes 1-6). Quantitative results from at least three independent mVP40 VLP budding assays in the presence of 0.3μM or 0.1μM concentration of FC-10696 revealed an average of 91% and 82% inhibition of mVP40 VLP egress, respectively compared to DMSO alone (Table 1).

**Table 1.**
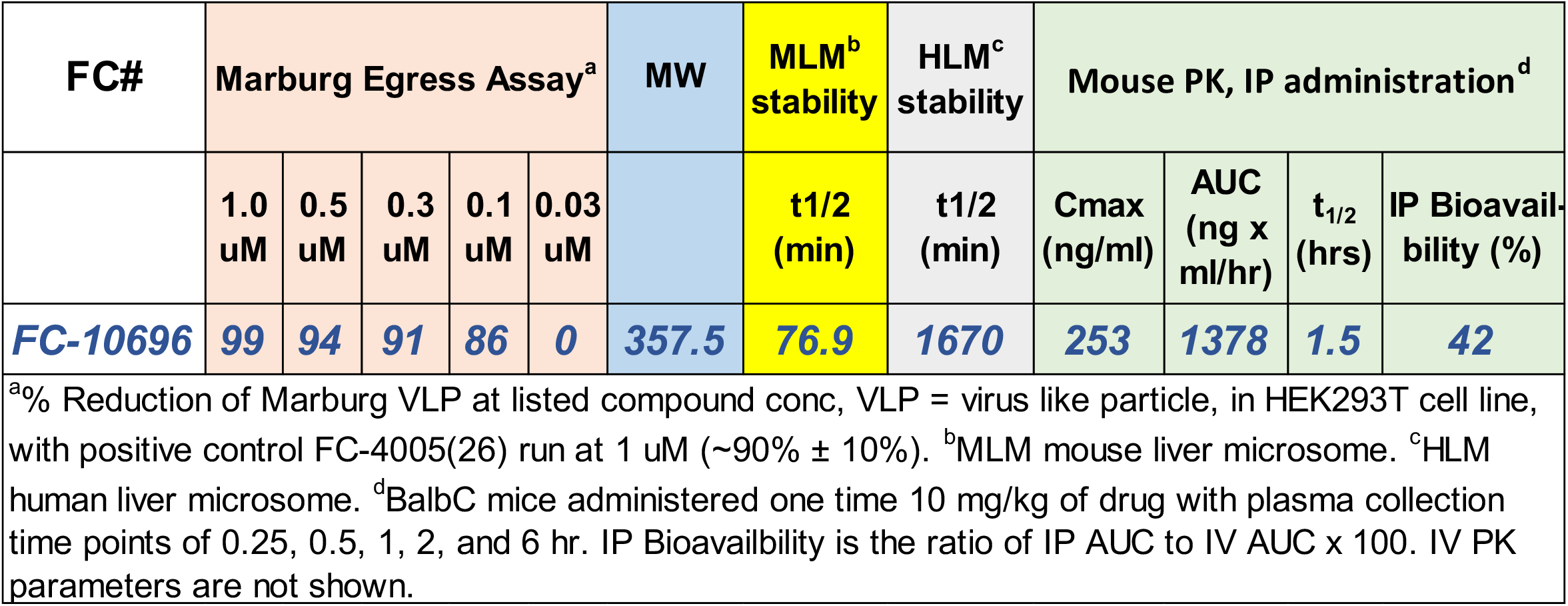
In vitro Marburg Egress, ADME and PK data

**Fig. 1.**
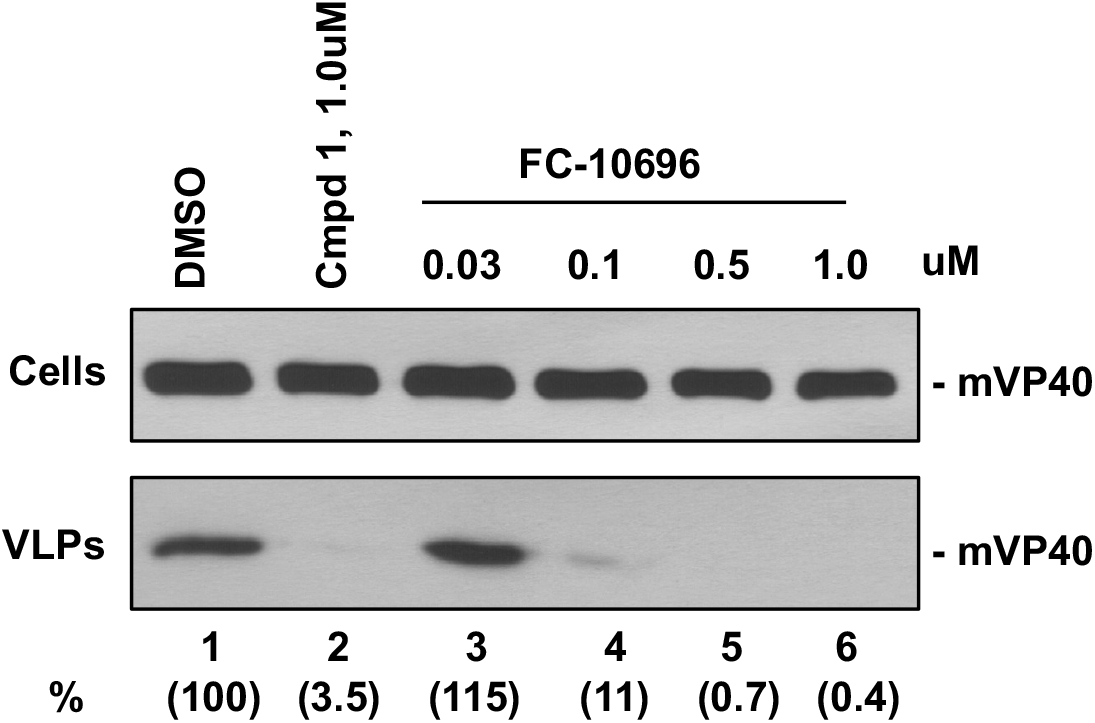
FC-10696 inhibits budding of mVP40 VLPs in a dose-dependent manner. HEK293T cells were transfected with mVP40 in the presence of DMSO alone, compound 1, or the indicated concentrations of compound FC-10696. mVP40 was detected in cell lysates and VLPs by Western blotting, and mVP40 levels in VLPs were quantified using NIH Image-J software (shown in parentheses).

We evaluated compound FC-10696 in a single dose mouse PK experiment under IP administration (Table 1). FC-10696 displayed excellent blood levels and good metabolic stability (Table 1). Indeed, FC-10696 was deemed suitable for live virus and mouse efficacy experiments, as it showed good stability to mouse and human liver microsomes and did not inhibit cytochrome P450 3A4 at concentrations up to 33uM, thus showing low risk for drug/drug interactions. In sum, FC-10696 had overall superior ADME and PK properties, as well as robust anti-budding activity in an mVP40 VLP budding assay.

### Compound FC-10696 inhibits egress and spread of live MARV in cell culture

To assess safety of FC-10696, we first assessed its cytotoxicity in a HeLa cell line, routinely used in anti-filoviral screens, and human monocyte-derived macrophages (MDMs), the initial targets of filovirus infection in the host (31, 32). Cells treated with 2-fold dilutions of FC-10696 or DMSO as a control were assessed for the number of metabolically active cells after 48 or 72 h. The CC_50_ value, the concentration that reduced the cell viability by 50% when compared to untreated control, showed that the compound was more cytotoxic to HeLa cells than to MDMs (Table 2 and Fig. 2A). Assessment of efficacy revealed that FC-10696 efficiently inhibited live MARV replication and egress in both cell types (Figs. 2B-D). Notably, the half maximal inhibitor concentration, IC_50_, value in the virion egress assay was at nanomolar concentrations, and the selectivity index, SI_50_, calculated as CC_50_/IC_50_, was ≥10, signifying antiviral potency and selectivity of these compounds against this virus.

**Table 2.**
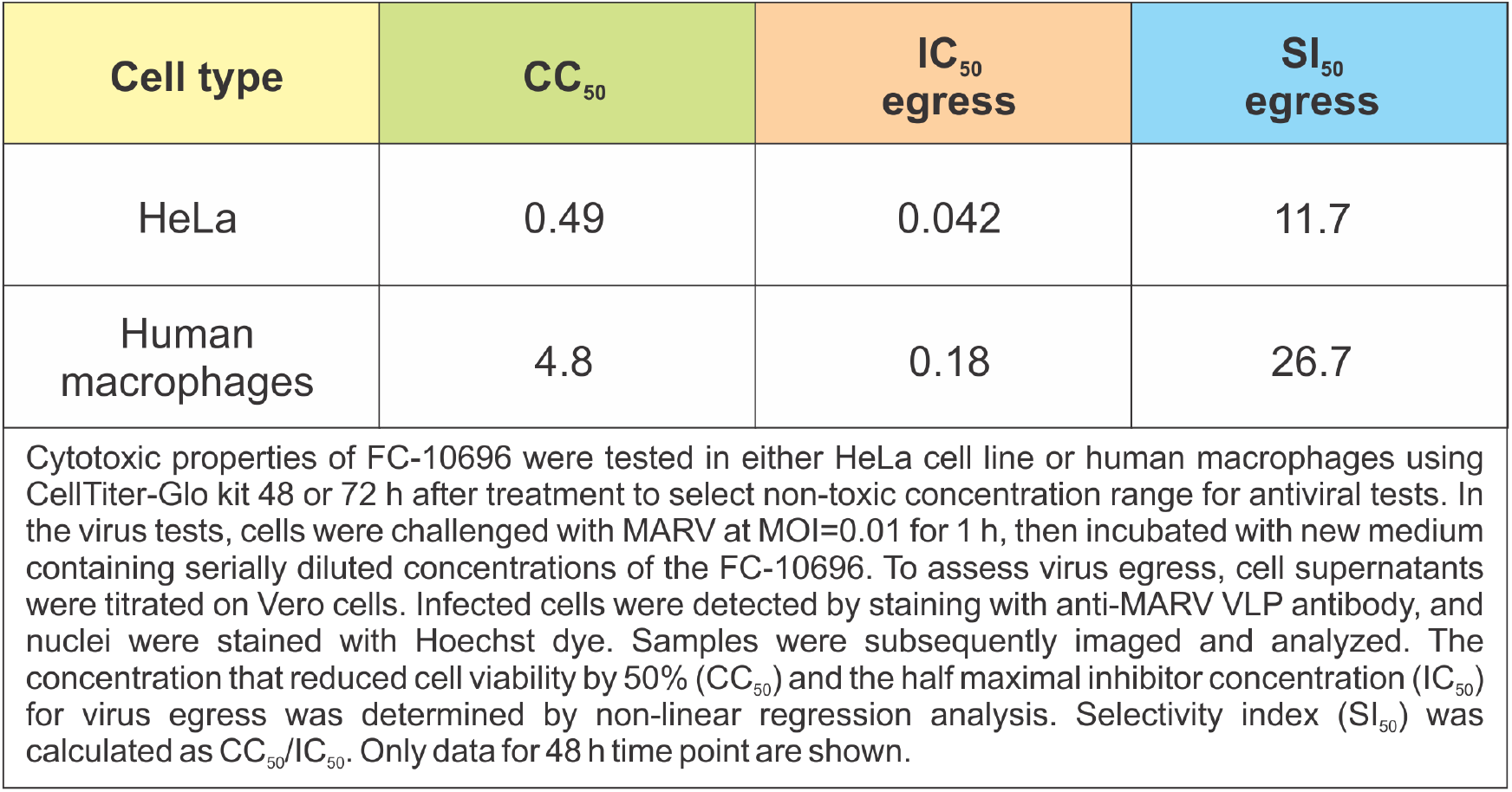
FC-10696 IC_50_ and S1_50_ values (in μm) for live MARV egress at 48 h.

| Cell type | CC<sub>50</sub> | IC<sub>50</sub><br>egress | SI<sub>50</sub><br>egress |
| --- | --- | --- | --- |
| HeLa | 0.49 | 0.042 | 11.7 |
| Human<br>macrophages | 4.8 | 0.18 | 26.7 |

Cytotoxic properties of FC-10696 were tested in either HeLa cell line or human macrophages using CellTiter-Glo kit 48 or 72 h after treatment to select non-toxic concentration range for antiviral tests. In the virus tests, cells were challenged with MARV at MOI=0.01 for 1 h, then incubated with new medium containing serially diluted concentrations of the FC-10696. To assess virus egress, cell supernatants were titrated on Vero cells. Infected cells were detected by staining with anti-MARV VLP antibody, and nuclei were stained with Hoechst dye. Samples were subsequently imaged and analyzed. The concentration that reduced cell viability by 50% (CC<sub>50</sub>) and the half maximal inhibitor concentration (IC<sub>50</sub>) for virus egress was determined by non-linear regression analysis. Selectivity index (SI<sub>50</sub>) was calculated as CC<sub>50</sub>/IC<sub>50</sub>. Only data for 48 h time point are shown.

**Fig. 2.**
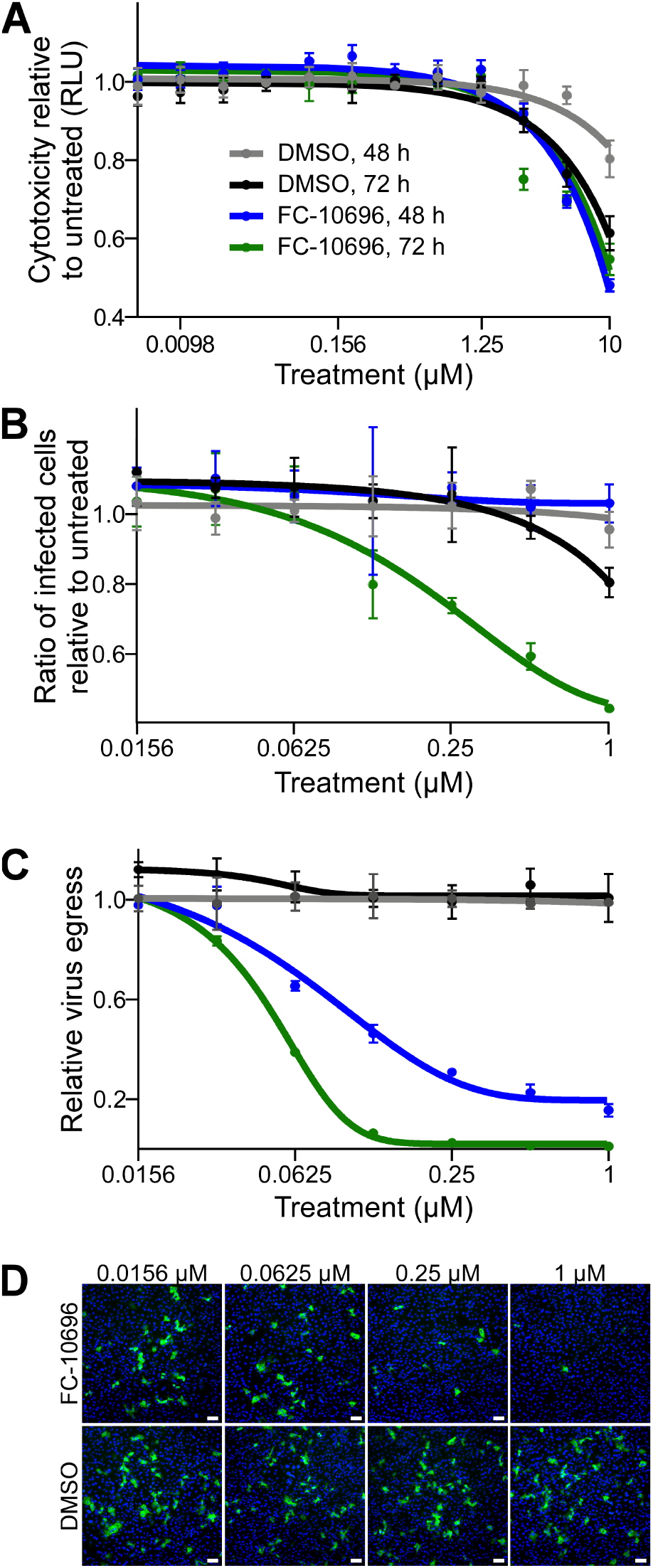
FC-10696 inhibits egress and spread of live MARV from infected human MDMs. **(A)** MDMs were left untreated or treated with FC-10696 at 11 2-fold serially diluted concentrations or DMSO for 48 or 72 h in triplicate. The number of metabolically active cells was determined using a CellTiter-Glo kit. The relative light unit (RLUs) value for each concentration is an average ± standard deviation of 3 replicates. **(B)** MDMs were challenged with MARV for 1 h, then washed and incubated with new medium containing 7 2-fold serially diluted concentration of the compounds, equal concentrations of DMSO, or no treatment, for 48 or 72 h. Subsequently, cells were stained with anti-VLP antibody and Hoechst dye, and photographed. Numbers of nuclei and infected cells were counted using CellProfiler software. The relative infection efficiencies, determined by dividing the number of infected cells by the number of nuclei, are reported relative to the infection efficiency in untreated cells and are averages ± standard deviations of 3 replicates. **(C)** To assess virus egress from FC-10696-treated MDMs, cell supernatants were titrated on Vero cells for 24 h. The samples were then treated with anti-VLP antibody and Hoechst dye. Samples were imaged and analyzed as above. **(D)** Supernatants of MDMs challenged with MARV and treated as indicated for 72 h were titrated on Vero cells. Samples were treated with anti-VLP antibody to detect infected cells (green) and Hoechst dye to detect nuclei (blue) and imaged using a Nikon imaging system. The bar in each image is 100 μm.

### Compound FC-10696 shows efficacy in a mouse challenge model of MARV infection

To evaluate antiviral potential of FC-10696 compound *in vivo*, BALB/cJ mice challenged intraperitoneally (IP) with 1,000 plaque-forming units (PFU) of mouse-adapted MARV were IP-dosed with a formulation containing the compound twice daily (BID) for 10 consecutive days, starting 6 h post-challenge. BALB/cJ mice are highly susceptible to infection with mouse-adapted MARV, developing disease symptoms and high viremia approximately three days post-infection, and succumbing to the disease by day 6 (33). We found that treatment with 20 mg/kg delayed the onset of mortality (p=0.0182; Fig. 3A), weight loss (Fig. 3B), and virus load in serum (p=0.0255; Fig. 3D). Importantly, the compound was well-tolerated in mice at the 20 mg/kg, showing that animals developed only transient treatment-associated toxicity (Fig. 4). Our results demonstrate the proof-of-concept *in vivo* activity for FC-10696 compound against MARV infection.

**Fig. 3.**
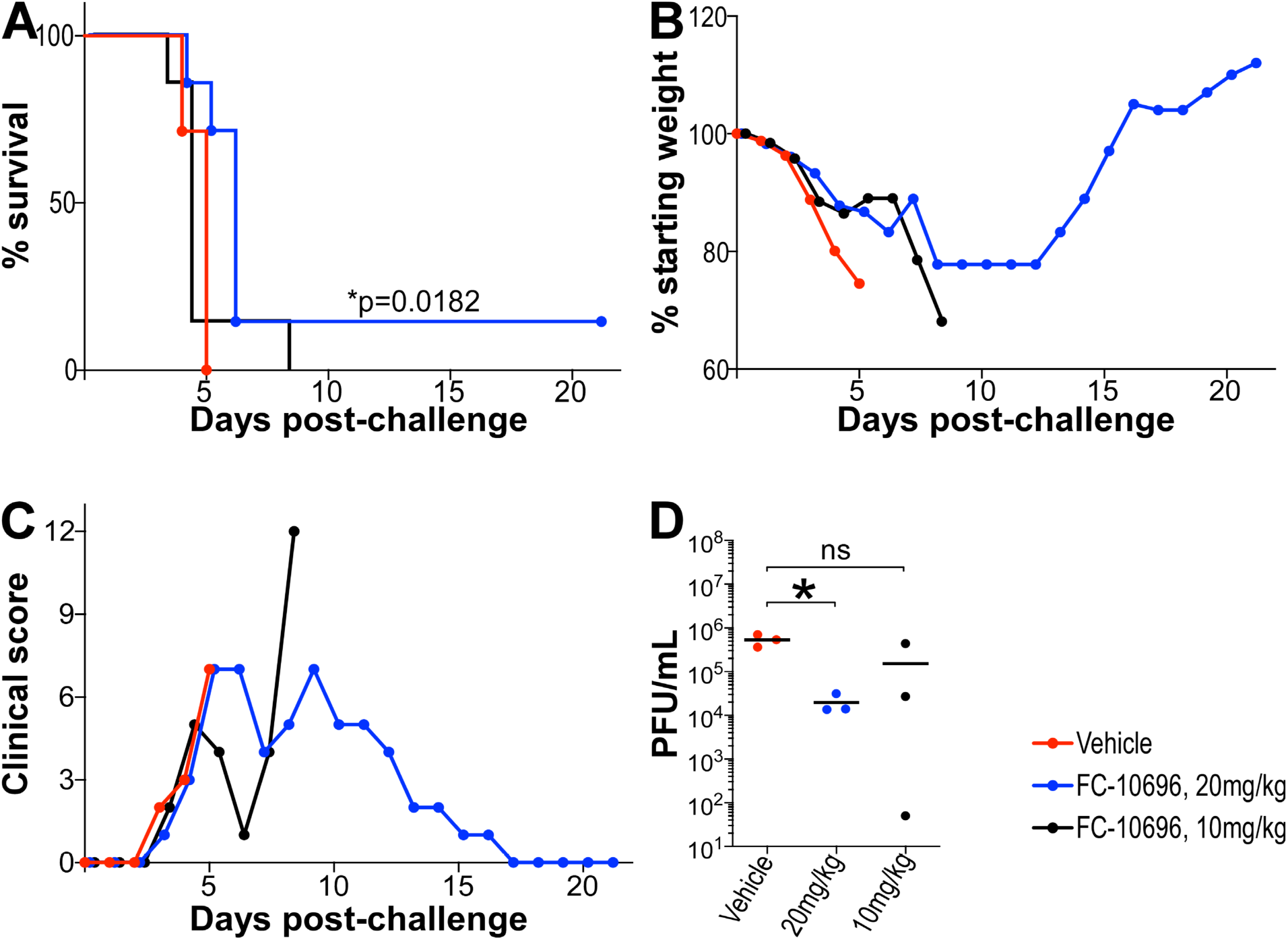
In vivo efficacy of FC-10696 in a mouse model of MARV disease. Three groups of 10 4-week old female BALB/cJ mice were challenged with 1,000 PFUs of mouse-adapted MARV. IP dosing by vehicle or FC-10696 at either 20 or 10 mg/kg started 6 h post-challenge and continued BID for 10 consecutive days. Animals were observed daily for mortality **(A)**, weight loss **(B)**, and clinical signs of disease **(C)** for 21 days post-infection. Clinical scores for each group were recorded as a sum of all observations in the group, and if a score of ≥12 was recorded for an individual animal, it was considered terminally ill and euthanized. On day 3 post-challenge, 3 animals/group were euthanized to collect serum for virus load assessment by the neutral red plaque assay **(D)**. The remaining 7 mice/group were used to determine animal survival. Viral burden was analyzed using a Student *t*-test or one-way ANOVA Tukey’s test, and survival analysis was performed using a Log-rank (Mantel-Cox) test, with p≤0.05 considered significant in all analyses.

**Fig. 4.**
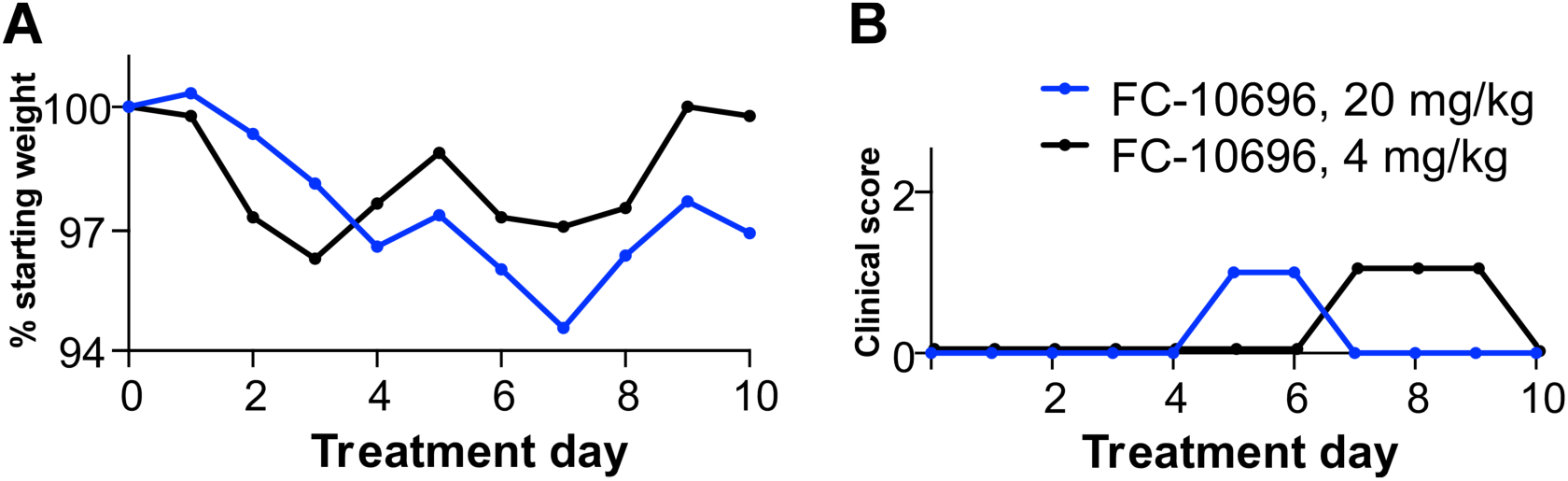
*In vivo* toxicity data for FC-10696. FC-10696 was resuspended in 30%PEG400/ 2%DMSO/14%kleptose HPB parenteral grade formulation at two different concentrations, 20 and 4 mg/kg, and administered to groups of five 4-week old female BALB/cJ mice BID via the IP route for a period of 10 days. Animals were monitored daily for signs of treatment-associated toxicity: weight loss **(A)**; rough hair coat, discharge from eyes and nose, diarrhea, decreased food intake and activity **(B)**; and mortality. Clinical scores for each group were recorded as a sum of all observations in the group.

## Discussion

The identification of host-oriented antiviral compounds represents a promising strategy to develop effective, broad-spectrum therapeutics capable of targeting a wide array of emerging pathogens, which may lead to a paradigm shift in the search for new antivirals (25–28, 34–43). MARV and other filoviruses continue to emerge and cause outbreaks of severe hemorrhagic fever largely originating in Africa, but with the potential to spread globally as observed with the EBOV outbreak in 2014-2015. Here, we report on our continued efforts to develop compounds targeting viral PPxY-mediated host interactions as a novel class of antiviral therapeutics. We demonstrate the antiviral efficacy of novel compound FC-10696 in an in vitro and in vivo model of MARV infection.

Extensive SAR led to the identification of compound FC-10696, which possesses excellent overall ADME and PK properties. FC-10696 exhibited potent activity at the BSL-2 level in blocking egress of mVP40 VLPs at low nanomolar concentrations in repeated experiments, and in disrupting a PPxY/WW-domain mediated interaction between mVP40 and human E3 ubiquitin ligase Nedd4 as determined using a bimolecular complementation (BiMC) approach (data not shown). FC-10696 was then moved into the BSL-4 laboratory where it was assessed for cytotoxicity in both Hela cells and hMDMs, as well as in BALB/c mice. After obtaining a satisfactory cytotoxicity profile and CC_50_ values, we went on to demonstrate that FC-10696 significantly inhibited MARV egress and spread from primary human macrophages compared to controls. Most intriguingly, we went on to show that treatment of mice with 20 mg/kg twice daily for 10 days delayed the onset of mortality (p=0.0182), weight loss, and virus load in serum (p=0.0255), thus providing strong support for this class of compounds for further development into potent antivirals against filoviruses and possibly other viruses that utilize PPxY L-domain motifs for productive infection. These findings represent the first proof-of-concept in vivo activity for our lead host-oriented PPxY inhibitor.

Studies are currently underway to assess compound FC-10696 and similar analogs for antiviral potency against related PPxY-containing viruses including EBOV and LASV. Indeed, preliminary findings indicate that FC-10696 can block egress and spread of both eVP40 VLPs and authentic EBOV in vitro; albeit less efficiently than MARV. This may be due to the presence of a second overlapping PTAP L-domain motif with the EBOV VP40 protein that in addition to the PPxY motif, can also function in promoting efficient egress and spread of EBOV, which is in contrast to the single isolated PPxY motif present in the MARV VP40 protein (24). As such, one could envision the use of a PPxY-mediated inhibitor such as FC-10696 in combination with a PTAP-mediated inhibitor, or a viral entry inhibitor for example as part of an antiviral cocktail strategy targeting multiple stages of the virus lifecycle for maximal effect (27, 30, 44–46). Our long term goal is to develop these compounds alone or in combination primarily for individuals in high risk situations including those in the military, health care workers, and first-line responders during an outbreak or epidemic. Additional studies are still needed to precisely identify the mechanism of action and drug interaction site, as well as additional studies to assess cytotoxicity and potential effects on the host in a wider array of cell types.

## Materials and Methods

### Cells, Plasmids, and Virus Strain

HEK293T, HeLa, and Vero cells were maintained in Dulbecco’s modified Eagle’s medium (DMEM) supplemented with 10% fetal bovine serum, penicillin (100U/ml)/streptomycin (100μg/ml), and the cells were grown at 37°C in a humidified 5% CO_2_ incubator. Flag-tagged mVP40 plasmid was kindly provided by S. Becker (Institut für Virologie, Marburg, Germany). Monocyte-derived human macrophages (MDMs) were isolated as described previously (47, 48). Peripheral blood was collected from healthy donors according to University of Texas Health approved IRB protocol 20180013HU. Heparinized blood was over-layed onto a Ficoll-Paque cushion (GE Heathcare, Uppsala, Sweden) to isolate peripheral blood mononuclear cells (PBMCs). PBMCs were cultured in suspension in RPMI medium supplemented with 20% autologous serum for 6 days at 37°C in a humidified 5% CO_2_ incubator to differentiate monocytes into macrophages. All experiments with live MARV were performed in the biosafety level 4 (BSL-4) laboratory at the Texas Biomedical Research Institute (TBRI, San Antonio, TX). MARV strain Musoke (NCBI accession number NC_001608) was obtained from the virus repository at the Texas Biomedical Research Institute. Mouse-adapted MARV strain Angola (NCBI accession number KM_261523) was generously provided by the National Microbiology Laboratory, Public Health Agency of Canada. Virus stocks were generated and characterized as described previously (33, 49).

### ADME and PK data

Human and mouse liver microsome stability studies and pharmacokinetic studies in mice were performed at Alliance Pharma, Inc. (17 Lee Boulevard, Malven, PA 19355). PK parameters from the PK study were calculated by M. Saporito.

### Human and mouse liver microsome stability

FC-10696, at a concentration of 0.5 uM was incubated with 0.5 mg/mL of liver microsomes (mouse or human) and an NADPH-regenerating system (cofactor solution) in potassium phosphate buffer (pH 7.4). At 0, 5, 15, 30, and 45 minutes, an aliquot was taken, and reactions were quenched with an acetonitrile solution containing an internal standard. Midazolam was run as a reference standard. Additionally, controls were measured that do not contain the cofactor solution. Following completion of the experiment, samples were analyzed by LC-MS/MS. Results were reported as peak area ratios of each analyte to internal standard. The intrinsic clearance {CL_int_) was determined from the first-order elimination constant by nonlinear regression.

### Pharmacokinetics in mice

A single-dose study was conducted in adult BalbC male mice of weight range 20-26 g each. Groups of six animals were administered an intravenous 2 mg/Kg (IV) dose or an intraperitoneal 10 mg/Kg (IP) dose of FC-10696 both administered as a soluble 5% DMSO/20% Kleptose aqueous formulation. Plasma samples (n= 3 per timepoint) were collected from study animals at 5, 15, 30 min; 1, 2, 6 hrs for the IV dose and 15, 30 min; 1, 2, 6 hrs for the IP dose. Collected samples were analyzed by LC-MS/MS. PK parameters were calculated using Prism Graphpad.

### VLP Budding Assays

MARV VP40 VLP budding assays in HEK293T cells were described previously (2). For VLP budding, HEK293T cells were transfected with 0.5μg of mVP40, and cells were treated with DMSO alone or the indicated concentration of inhibitor for 24 hours post transfection. The mVP40 protein in cell extracts and VLPs was detected by SDS-PAGE and Western blotting and quantified using NIH Image-J software. Anti-flag monoclonal antibody was used to detect flag-tagged mVP40.

### MARV Egress and Spread Assays

HeLa cells or MDMs were plated into wells of a 96-well plate at 2×10^4^ or 5×10^4^ cells/well, respectively, to determine cytotoxic and antiviral properties of FC-10696. All treatments were performed in triplicate. In cytotoxicity assays, cells were left untreated or treated with the compound at eleven 2-fold serially diluted concentrations or DMSO (solvent) for 48 or 72 h. The number of metabolically active cells was determined using a CellTiter-Glo kit. The concentration that reduced the cell viability by 50% when compared to untreated control, CC_50_ value, at each time point was determined using non-linear regression analysis using GraphPad 8 software to select a non-toxic concentration range for antiviral tests.

In virus tests, HeLa cells or MDMs were challenged with MARV at multiplicity of infection (MOI) of 0.01 for 1 h to allow binding, then washed and incubated with new medium containing seven 2-fold serially diluted concentration of the compounds, equal concentrations of DMSO, or no treatment, for 48 or 72 h. To assess virus egress, cell supernatants were titrated on Vero cells for 24 h. Infected cells were detected by treatment with MARV VLP antibody (IBT Bioservices, Rockville, MD), and nuclei by staining with Hoechst dye (Thermo Fisher Scientific, Waltham, MA). Samples were photographed using a Nikon automated system (Nikon, Tokyo, Japan) and analyzed by CellProfiler software (Broad Institute, Cambridge, MA) to quantify virus spread and egress. Infection efficiency in treated samples was determined as a ratio of infected cells and nuclei and reported relative to mock. The half maximal inhibitor concentration, IC_50_, value for virus spread and egress for each time point was determined by non-linear regression analysis. The selectivity index, SI_50_, determined as CC_50_/IC_50_, was used to assess antiviral potential of the compounds.

### Animals

Wild-type 4-week old female BALB/cJ mice were obtained from The Jackson Laboratory (Bar Harbor, ME). The mouse studies were conducted in strict adherence to the Animal Welfare Act and the Guide for the Care and Use of Laboratory Animals of the National Institutes of Health (NIH). The TBRI animal assurance welfare number is D16-00048 (A3082-01) under file with the NIH. All mouse procedures were approved by the TBRI Institutional Animal Care and Use Committee (IACUC) which oversees the administration of the IACUC protocols. The mouse studies were performed as outlined in the IACUC protocol #1708MU.

### *In Vivo* Efficacy Studies

To assess toxicity of FC-10696 treatment in mice, the compound was resuspended in 30%PEG400/ 2%DMSO/14%kleptose HPB parenteral grade (Roquette, Lestrem, France) formulation at two different concentrations, 20 and 4 mg/kg, and administered to groups of five 4-week old female BALB/cJ mice twice daily (BID) via the intraperitoneal route (IP) for a period of 10 days. Animals were monitored daily for signs of treatment-associated toxicity: weight loss, rough hair coat, discharge from eyes and nose, diarrhea, decreased food intake and activity, and mortality. Clinical scores for each group were recorded as a sum of all observations in the group.

To assess antiviral potential of FC-10696 treatment in a mouse model of MARV disease, three groups of ten 4-week old female BALB/cJ mice were challenged with 1,000 plaque-forming units (PFUs, as determined on Vero cells) of mouse-adapted MARV by the IP route. IP dosing by vehicle or FC-10696 at either 20 or 10 mg/kg started 6 h post-challenge and continued BID for 10 consecutive days. Animals were observed at least twice daily for signs of viral disease (ruffled hair coat, hunch back, inappetence, weight loss, and decreased movement) and mortality for 21 days post-challenge. Group clinical scores were recorded as the sum of all clinical observations for the group. If a clinical score of ≥12 was recorded for an animal, it was considered terminally ill and euthanized. Three animals from each group were euthanized on day 3 post-challenge to collect blood to determine virus titer by a plaque assay. The remaining 7 mice/group were used to determine animal survival.

## Acknowledgments

Funding was provided in part by National Institutes of Health grants AI138052 and AI138630 to RNH, AI129890 to BDF, T32-AI070077 to ASM, and an Innovator Award from The Wellcome Trust to MSS. The funders had no role in study design, data collection and analysis, decision to publish, or preparation of the manuscript.

